# Opponent regulation of striatal output pathways by dopamine and serotonin

**DOI:** 10.1101/2025.04.17.649431

**Authors:** Daniel F. Cardozo Pinto, Michaela Y. Guo, Wade Morishita, Matthew B. Pomrenze, Neir Eshel, Robert C. Malenka

**Author notes:** Department of Molecular and Cellular Biology, Harvard University, Cambridge, MA 02138, USA. These authors contributed equally (Daniel F. Cardozo Pinto, Michaela Y. Guo).

## Abstract

Classic theories propose opponent functions for striatal dopamine (DA) and serotonin (5-hydroxytryptamine; 5HT), with DA promoting approach and 5HT promoting patience or avoidance. How these neuromodulators regulate downstream circuits to achieve such antagonistic effects remains mysterious. Here, we mapped striatal 5HT receptor expression and recorded from genetically-identified striatal neurons to demonstrate that DA and 5HT selectively activate distinct populations of striatal neurons to exert opponent control over striatal output.

## MAIN

Striatal DA release is crucial for animals to learn from and seek rewards^1^. Longstanding theories have proposed that 5HT works against DA by promoting behavioral suppression or driving aversive learning^2–7^. These influential notions of opponency are supported by pharmacology experiments showing that 5HT-releasing drugs blunt the DA-driven rewarding effects of drugs of abuse^8–11^, as well as by studies showing that 5HT activity can suppress reward-seeking behavior^12–16^. Consistent with an oppositional relationship between striatal DA and 5HT, we recently found that rewards trigger opposite striatal DA and 5HT responses in a single animal and that both of these inverse signals are required for a stimulus to be maximally reinforcing^17^. However, it remains unclear how opponent DA and 5HT signals regulate striatal activity to exert opposing effects on behavior.

In the striatum, DA works at least in part by differentially modulating excitability and plasticity in two primary projection neuron classes. DA is thought to increase activity in medium spiny neurons expressing the Gs-coupled D1 DA receptor (D1-MSNs) and decrease activity in those expressing the Gi-coupled D2 receptor (D2-MSNs)^18,19^. However, we do not have a similar understanding of the effects of 5HT on striatal circuits. Owing to the limited spatial and cell-type resolution of transcriptomic^20^ and autoradiographic^21^ studies, respectively, which 5HT receptors are expressed across striatal subregions and cell-types remains poorly understood.

To address this knowledge gap, we mined a single-cell RNA-sequencing dataset of the mouse striatum^22^ to identify which 5HT receptors are expressed in MSNs. We found that the vast majority (>90%) of MSNs express 5HT receptors, including members of the Gi-, Gs-, and Gq-coupled 5HT receptor families but not the ionotropic receptor 5HTR3 (**Extended Data Fig. 1a-c**). Of 14 5HT receptor genes, 7 are widely expressed in MSNs (*Htr1b*, *Htr1d*, *Htr1f*, *Htr2a*, *Htr2c*, *Htr4*, *Htr6;* **Extended Data Fig. 1d,e**), and one more is expressed in a unique minority MSN population in rodents^22,23^ and humans^24^ (*Htr7*). Some 5HT receptor genes showed differential expression between striatal patch and matrix compartments and many were correlated with expression of D1- or D2- MSN marker genes (**Extended Data Fig. 1f,g)**. Together, these results suggest that 5HT receptor expression in striatal MSNs is nearly ubiquitous, restricted primarily to 8 receptor genes, and varied across cell-types and subregions.

To better understand striatal 5HT receptor organization, we used fluorescence *in situ* hybridization to map expression of the subset of genes identified above. In each experiment, we labeled transcripts for the D1 receptor *(Drd1a*), the D2 receptor (*Drd2*), and one 5HT receptor gene, allowing us to identify putative D1- and D2- MSNs (pD1- and pD2-MSNs, respectively) and measure spatial location and receptor expression individually for each cell. Control samples labeled for the MSN marker gene *Ppp1r1b* validated that this approach accurately identifies MSNs which could be divided into pD1 and pD2 subpopulations based on their DA receptor profiles (see Methods and **Extended Data Fig. 2a-e**).

Expression of every 5HT receptor gene we examined varied across cell-types and subregions (**Fig. 1a-h**). pD1-MSNs preferentially expressed one Gi-coupled receptor (*Htr1f*) and one Gs-coupled receptor (*Htr4*), suggesting 5HT could have a relatively balanced net effect on their activity (**Fig. 1c,f**). By contrast, pD2-MSNs preferentially expressed one Gi-coupled (*Htr1d*) and four Gq- (*Htr2a*, *Htr2c*) or Gs- (*Htr6, Htr7*) coupled receptors, suggesting 5HT release is likely to increase activity in these cells (**Fig. 1b,d,e,g,h** and **Extended Data Fig 3a-d**). Spatially, individual 5HT receptor genes were preferentially expressed in dorsal (*Htr7*), ventral (*Htr1f*, *Htr4*), medial (*Htr2c*), or lateral striatum (*Htr2a*), and some were enriched (*Htr2a*, *Htr7*) or depleted (*Htr1b*) in irregular patches consistent with striosome and matrix organization, respectively (**Fig. 1i**).

**Fig. 1:**
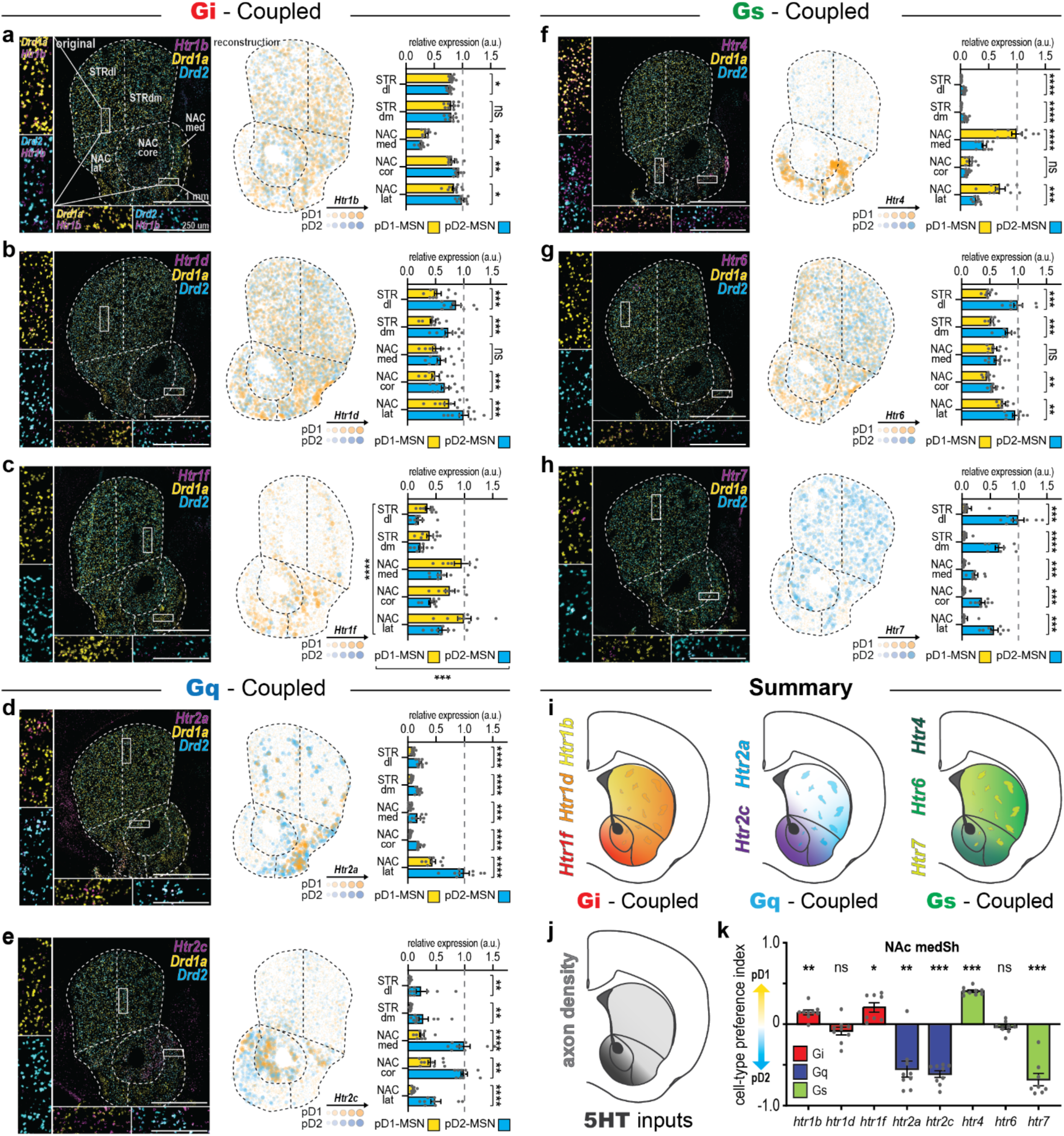
Differential 5HT receptor expression profiles between pD1- and pD2- MSNs. **a,** Left, fluorescence *in situ* hybridization labeling of *Drd1a, Drd2, and Htr1b* in the striatum. Center, reconstruction of the relative expression of *Htr1b* in the striatum. Each pMSN is represented by a dot, dots are colored by cell-type, and the size and opacity of each dot scales with the relative expression of *Htr1b* in that cell. Right, quantification of the relative expression of *Htr1b* across striatal subregions and cell-types. **b,** Same as **a**, but for *Htr1d*. **c,** Same as **a**, but for *Htr1f*. **d,** Same as **a**, but for *Htr2a*. **e,** Same as **a**, but for *Htr2c*. **f,** Same as **a**, but for *Htr4*. **g,** Same as **a**, but for *Htr6*. **h,** Same as **a**, but for *Htr7*. **i-j,** Schematics summarizing relative 5HT receptor expression (**i**) and 5HT axon density (**j**) across striatal subregions. **k,** Cell-type preference analysis for 5HT receptor expression in the NAc^medSh^. For this and all subsequent figures, data are shown as mean ± s.e.m., significance is denoted as *P < 0.05, **P < 0.01, ***P < 0.001, ****P < 0.0001, significance tests were two-tailed, and statistics are shown in Supplementary Table 1.

This spatial heterogeneity motivated us to ask which striatal subregions receive the most 5HT input. By analyzing labeled axons from dorsal raphe 5HT (DR^5HT^) neurons^17^ – the primary source of striatal 5HT input^25^ – we found that 5HT inputs were denser in the nucleus accumbens (NAc) compared to the dorsal striatum (consistent with reports in rats^26–28^ and primates^29,30^; **Extended Data Fig. 4a,b**), and densest in the posterior NAc medial shell (NAc^medSh^; **Fig. 1j** and **Extended Data Fig. 4c-g**). Examining cell-type differences in 5HT receptor expression in this region suggested that striatal pD1- and pD2-MSNs may be differentially regulated by 5HT, likely in a manner opposite to their modulation by DA (**Fig. 1k**).

To test this hypothesis, we performed acute slice recordings from genetically identified D1- and D2-MSNs in the NAc^medSh^ and measured their responses to bath application of DA or 5HT (**Fig. 2a**). We did this in cell-attached mode to prevent dialyzing the cell – which interferes with the intracellular signal transduction cascades recruited by G-protein coupled receptors^31,32^ – and in the presence of potassium channel blockers to induce spontaneous activity in normally quiescent MSNs, allowing us to detect both increases and decreases in MSN activity in response to DA or 5HT (**Fig. 2b,c**). In agreement with previous work^18,19,32^, we found that DA increased activity of D1-MSNs and decreased activity of D2-MSNs (**Fig. 2d-g** and **Extended Data Fig. 5a,b**). By contrast, bath application of 5HT did not significantly change D1-MSN activity, but it excited D2-MSNs (**Fig. 2h-k** and **Extended Data Fig. 5c,d**), consistent with these cell-types’ distinct 5HT receptor expression profiles. Thus, we found that DA and 5HT differentially regulate the activity of the two primary striatal cell-types (**Fig. 2 l-n**).

**Fig. 2:**
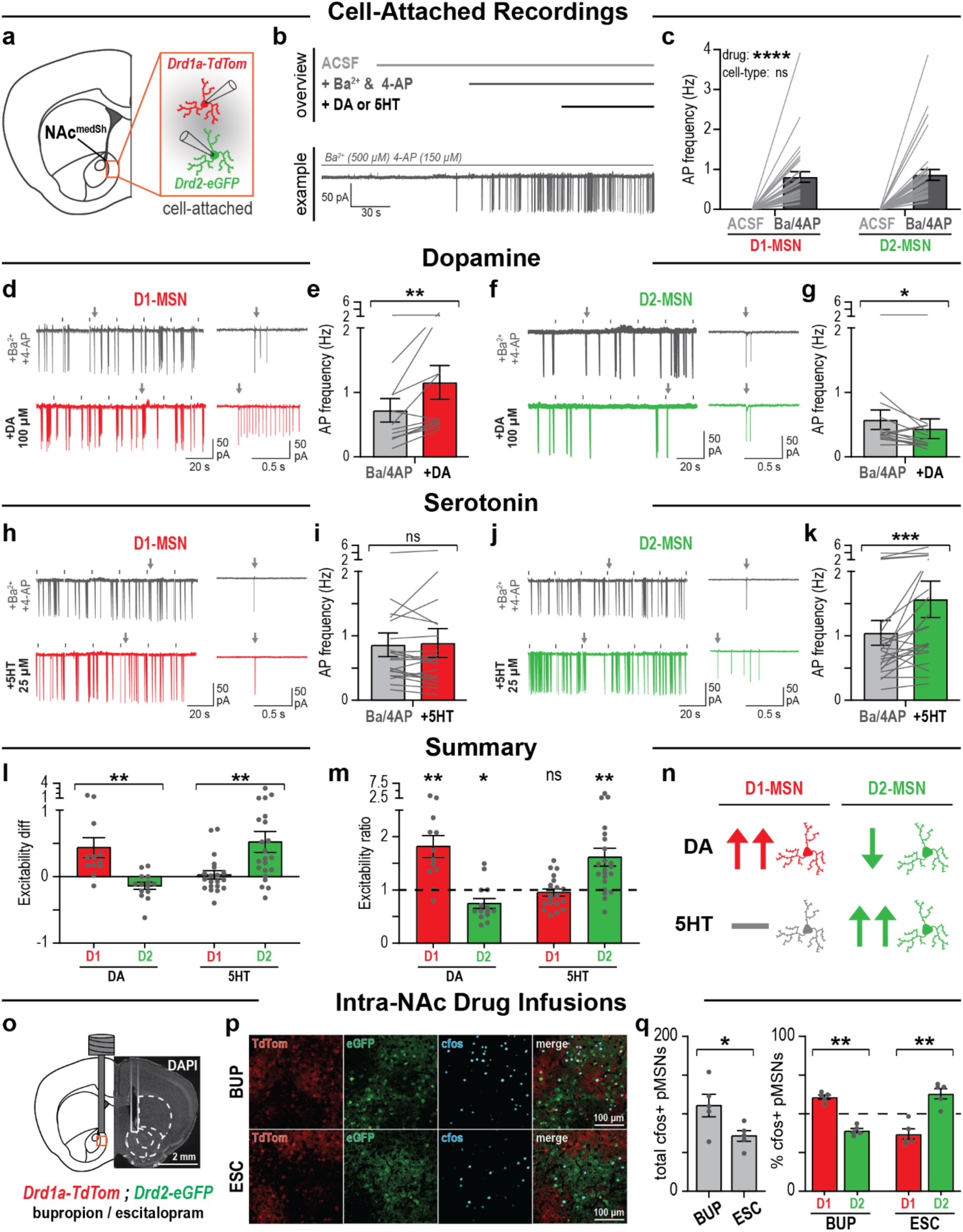
DA and 5HT differentially regulate activity of D1- and D2-MSNs. **a,** Genetic strategy for cell-attached recordings from D1- and D2-MSNs. **b,** Top, schematic of experimental design for cell-attached recordings. Bottom, example trace showing addition of Ba^2+^ and 4-AP induces spontaneous activity in an MSN. **c,** Spontaneous activity induced by Ba^2+^ and 4-AP did not differ between D1- and D2- MSNs. **d-e,** Example traces depicting D1- MSNs before (top) and after (bottom) addition of DA (**d**), and corresponding quantification of action potential frequency (**e**). **f-g**, same as **d-e** but for D2-MSNs. **h-i,** Example traces depicting D1-MSNs before (top) and after (bottom) addition of 5HT (**d**), and corresponding quantification of action potential frequency (**e**). **j-k**, same as **h-i** but for D2-MSNs. In **d**,**f**,**h**,**i**, tick marks indicate where calibration pulse artifacts were manually removed, and arrows indicate corresponding locations on the example traces (left) and insets (right). **l-m**, Relative changes (**l,** difference; **m**, ratio) in D1- and D2- MSN activity following application of DA and 5HT. **n,** Summary of cell-attached recording experiment results. **o,** Schematic and example image of cannula implantation site for intra-NAc infusion of bupropion or escitalopram in *Drd1a-TdTom;Drd2-eGFP* double transgenic mice. **p,** Example images of cfos expression in the NAc^pmSh^ of *Drd1a-TdTom;Drd2-eGFP* mice treated with bupropion (top) or escitalopram (bottom). **q**, Quantification of total (left) and relative (right) cfos expression in bupropion and escitalopram treated mice.

To test whether the differential regulation of D1- and D2-MSN activity by DA and 5HT that we observed in slice also occurs *in vivo*, we implanted *Drd1a-TdTomato;Drd2-eGFP* double transgenic mice with drug infusion cannulas above the NAc^medSh^ (**Fig. 2o**). We then selectively upregulated DA or 5HT activity in this region by locally infusing a DA reuptake inhibitor (bupropion) or a 5HT reuptake inhibitor (escitalopram) into the NAc and immunostained the brains for the immediate early gene product cfos; a molecular proxy for neural activation. Analyzing cfos expression in genetically-identified D1- and D2- MSNs revealed that bupropion-treated mice showed a greater number of cfos*+* MSNs overall compared to escitalopram-treated mice. Importantly, bupropion treatment preferentially induced cfos expression in D1 MSNs while escitalopram treatment preferentially induced cfos expression in D2 MSNs (**Fig. 2p,q**).

Together, our results suggest that DA and 5HT exert inverse effects over striatal outputs via differential regulation of D1- and D2-MSN activity in a manner congruent with these cell-types’ distinct 5HT receptor expression profiles. Preferential activation of D1- and D2- MSNs by DA and 5HT, respectively, may thus be one mechanism contributing to opponent control of reinforcement by striatal DA and 5HT^17^. Of note, this result provides a potential explanation for longstanding observations that 5HT2a^33–35^ and 5HT2c^36–39^ receptor activation dampen the reinforcing properties of addictive drugs because it suggests 5HT2 family receptors, which are strongly concentrated on D2-MSNs and depolarize striatal neurons^40,41^, counteract the inhibitory effect of drug-driven DA on these cells.

The effects we describe of DA and 5HT on striatal activity may be augmented by additional actions of these neuromodulators in the striatum, including modulating the strength of corticostriatal^42–44^ and thalamostriatal^45,46^ synapses, regulating lateral inhibition between D1 and D2 MSNs^47–49^, differentially filtering presynaptic inputs onto MSNs^46^, and/or differentially regulating striatal interneuron activity^50–52^. Our identification of cell-type specific expression of 5-HT receptor subtypes in the striatum should inspire further investigations into the mesolimbic 5HT system and how its interactions with DA shape a wide range of motivated behaviors.

## Supporting information

Supplemental Table 1 - statistics

## METHODS

### Mice

C57BL6/J (Jackson Laboratory #000664), *Drd1a-TdTomato* (Jackson Laboratory #016204), *Drd2-eGFP* (Jackson Laboratory #030255), and *Drd1a-TdTomato;Drd2-eGFP* double transgenic (cross between Jackson Laboratory #016204 and #030255) mice of both sexes (>5 weeks old) were used in this study. Mice were housed at ∼21°C with 30-70% humidity on a 12-hour light/dark cycle with ad-libitum access to food and water. All experimental procedures were approved by the Stanford University Administrative Panel on Laboratory Animal Care and the Administrative Panel on Biosafety.

### Stereotactic surgeries

Stereotactic surgeries were performed as previously described^17^. Mice were anesthetized with isoflurane (4-5% induction, 1-2% maintenance) and a stereotaxic frame (Kopf instruments) was used to target injections and implantations to the following structures (coordinates are in mm, with AP and ML relative to bregma and DV relative to the skull surface): DR, AP −4.6, ML 0, DV - 3.3; NAc^pmSh^, AP +1.0, ML +/−0.7, DV −4.3. Viral vectors (∼10^12^ gc/mL, from the Stanford Gene Vector and Virus core) were infused with a syringe pump (Harvard Apparatus) at a rate of 100-200 nL/min and allowed to diffuse for at least 5 min. Drug infusion guide cannulas were secured to the skull using screws (Antrin Miniature Specialties) and dental cement (Geristore). Viral incubation times were >3 weeks. Drug infusion guide cannulas placements were histologically verified post-hoc.

### Fluorescence *in situ* hybridization

Fluorescence in situ hybridization was performed as previously described^53^. Briefly, wild-type mice were deeply anesthetized with isofluorane and then decapitated. The brains were dissected and flash frozen in isopentane on dry ice. Coronal sections of the striatum (16 um thick, approximately from 0.74 to 1.70 mm anterior to bregma) were collected on a cryostat, and samples were prepared following the RNAscope Multiplex Fluorescent Reagent Kit v2 protocol from ACDBio, using 1 hr on ice cold PFA for fixation and 5 min treatment with Protease Plus for digestion. The following probes were used, all from ACDBio: *Drd1a* (#406491, #406491-C2), *Drd2* (#406501-C3), *Htr1b* (#315861-C2), *Htr1d* (#315871-C2), *Htr1f* (#315881-C2), *Htr2a* (#401291-C2), *Htr2c* (#401001-C2), *Htr4* (#408241), *Htr6* (#411161), *Htr7* (#401321-C2), and *Ppp1r1b* (#405901-C2). The *Drd1a* and *Drd2* probes were used for all experiments, while the third labeled gene varied across different experiments, and slides were coverslipped with Fluoromount-G containing DAPI (Southern Biotech, 0100-20). Thus, all samples were labeled for DAPI, *Drd1a*, *Drd2*, and one more gene that varied across experiments (either a 5HT receptor gene or the MSN marker gene *Ppp1r1b*).

Stitched images of the entire striatum in each hemisphere were acquired on a Keyence BZ-X800 microscope running BZ-X800 Viewer software using a 10x, 0.45 NA objective with a resolution of ∼0.755 um/pixel. Exposure times for the DAPI, *Drd1a*, and *Drd2* channels was kept constant across all experiments. Owing to differences in the expression level of the various 5HT receptor genes and the MSN marker gene *Ppp1r1b*, exposure times for imaging the third channel varied across experiments, but within an experiment all samples were imaged with identical settings and quantifications presented show only relative, not absolute, differences to account for the different exposure times used. Subsequent image analysis was carried out in ImageJ to perform cell-detection based on the DAPI channel and then measure signal in the other fluorescent channels separately for each detected cell. Specifically, each fluorescent channel was preprocessed as follows: DAPI channel, background subtraction (rolling ball, radius 50 pixels), top-hat filter (radius 12 pixels), sharpening, top hat filter (7 pixels); *Drd1a* and *Drd2* channels, background subtraction (rolling ball, radius 50 pixels), top-hat filter (radius 12 pixels); 5HT receptor gene or *Ppp1r1b* channel, background subtraction (rolling ball, radius 50 pixels). All preprocessed channels were then duplicated and one copy of each channel was binarized as follows: DAPI, thresholding using Li’s algorithm^54^, and then watershedding; *Drd1a* and *Drd2* channels, thresholding using Li’s algorithm^54^, erosion; 5HT receptor gene and *Ppp1r1b* channel, thresholding using the moments algorithm^55^. Samples were then manually annotated for striatal subregions based on DAPI signal and the Paxinos brain atlas^56^, and cell-detection was done by applying the ImageJ Analyze Particles function (size = 64-324, circularity = 0.25-1.00) to the binarized DAPI channel. Cell-ROIs were then overlayed onto the other preprocessed and binarized fluorescence channels to measure the mean brightness and percentage of area covered by fluorescence signal in each channel for each cell, and the coordinates and striatal subregion of each detected cell were also recorded.

Putative D1 and D2 MSNs were then identified on the basis of their *Drd1a* and *Drd2* signals. First, cells with low DA receptor expression (defined as <75% area covered by *Drd1a* or *Drd2* signal) were excluded. Then, a cell-type identity index was calculated ([*Drd1* area covered – *Drd2* area covered] / [*Drd1* area covered + *Drd2* area covered]) and cells with a score less than 0 were assigned the pD2-MSN label and cells with a score greater than 0 were assigned the pD1-MSN label. A very small minority of cells with a cell-type identity score of 0 (i.e. equal amounts of *Drd1a* and *Drd2* signal) were excluded from further analysis (see Extended Data Fig. 2e). These image preprocessing steps and the chosen cutoff values were validated using a dataset where the MSN marker gene *Ppp1r1b* was labeled instead of a 5HT receptor gene, as shown in Extended Data Fig 2. This control experiment confirmed that the image analysis pipeline described above accurately identified MSNs and the resulting population of cells showed a clear bimodal distribution of *Drd1a* and *Drd2* expression enabling us to assign pD1- and pD2- MSN identities accordingly.

5HT receptor expression was then compared across striatal subregions and cell-types as follows. 5HT receptor expression reconstruction maps in Fig 1 were generated by selecting an example hemisphere for each experiment and plotting each pMSN in that hemisphere as a dot at its corresponding coordinates in the original image. Each dot was then colored by cell-type and scaled in size and opacity by the signal in the 5HT receptor channel for that cell (percentage of each cell’s area covered by 5HT receptor gene signal, scaled separately for each 5HT receptor gene using the scale function in R with center=FALSE and scale=TRUE). The relative expression bar graphs in Fig 1 show the percentage of each cell’s area covered by 5HT receptor gene signal averaged within each subregion for each mouse and normalized to the highest expressing cell-type and subregion for that gene. Finally, the cell-type preference index graphs in Fig 1j and Extended Data Fig. 3 were generated by averaging the percentage of area covered by 5HT receptor signal across all the cells in each subregion for each mouse, and then calculating a cell-type preference index with the formula ([pD1-pD2]/[pD1+pD2]).

### RNAseq analysis

RNAseq data shown in Extended Data Fig 1a-g originated from Stanley et al^22^. We performed original analyses on this data beginning with the count matrix, which listed the number of reads detected of each gene for every cell that met the original paper’s quality control standards, and the metadata matrix which contained the discrete and continuous cell-type identities assigned to each cell in the count matrix. Cells from the olfactory tubercle and islands of Calleja (continuous subtypes “OT.ruffle”, “OT.flat”, and “ICj”) were excluded from analysis. For the remaining data, the expression level (as counts per million reads; cpm) and fraction of cells (as a percentage) expressing at least one read was calculated for each 5HT receptor gene and averaged within mice. For the patch vs matrix comparison, the same calculation was done separately for cells in each striatal compartment (patch: continuous subtypes “Htr7.vMedPat”, “Pat.Dorsolateral_CPu”, “Pat.Ventromedial_CPu”; matrix: continuous subtypes ““Mat.Dorsolateral_CPu”, “Mat.Ventromedial_CPu”). Finally, gene expression correlation analysis was done by calculating the Spearman correlation between scaled log expression levels for D1 and D2 marker genes and 5HT receptor genes for all cells pooled across mice.

### Histology and Immunohistochemistry

Histology and immunohistochemistry were performed as previously described^17^. Mice were transcardially perfused with 4% (w/v) paraformaldehyde (PFA) in phosphate-buffered saline (PBS) and the brains were postfixed in PFA overnight before being sectioned coronally to a thickness of 50 um on a vibratome. Sections were then washed three times for 10 min in PBS, blocked and permeabilized for 1 hr in a solution of 10% normal goat serum, 0.2% bovine serum albumin, and 0.5% triton-X, washed once for 5 min in PBS, incubated with primary antibodies in carrier solution (1% normal goat serum, 0.2% bovine serum albumin, and 0.5% triton-X) overnight on a shaker at room temperature, washed four times for 10 min, incubated for 2 hr with secondary antibodies in carrier solution, and finally washed four times for 10 min in PBS before being mounted on microscope slides and coverslipped with Fluoromount-G containing DAPI (Southern Biotech #0100-20). For axon tracing experiments, sections were immunostained for EYFP (primary: chicken anti-GFP, Aves #GFP-1020; secondary: goat anti-chicken 488, Invitrogen #A-11039) and/or mCherry (primary: rat anti-mCherry, Invitrogen #M11217; secondary: goat anti-rat 594, Invitrogen #11007). For immediate early gene experiments, sections were immunostained for cfos (primary: rabbit anti-cfos, Sysy #226008; secondary: goat anti-rabbit 647, Invitrogen # A-21244). Primary antibodies were used at a concentration of 1:1000 and secondary antibodies were used at a concentration of 1:750.

### Axon tracing

The axon tracing data presented in Extended Data Fig. 3a-e are a new analysis of data that were collected previously^17^. For the DR^5HT^ presynaptic bouton labeling experiments in Extended Data Fig. 3f-g, SERT-Cre mice were injected in the DR with 1 uL of AAV-DJ-hSyn-GFP-2A-synaptophysin-mRuby, immunostained for EYFP and mCherry, and example images were captured on a Nikon A1 confocal microscope.

### Intra-NAc drug infusions and cfos screen

Intra-NAc drug infusion experiments were performed on *Drd1a-TdTomato;Drd2-eGFP* double transgenic mice. Guide cannulas (Protech International) were surgically implanted unilaterally 1 mm above the NAc^pmSh^ (i.e. −3.3 mm DV) such that the tip of the infusion cannula (Protech International), which protruded 1 mm past the tip of the guide cannula, would reach the target coordinate on the day of the experiment. After recovery from surgery (>1 week), mice received 1-2 days of habituation to handling, to the open field behavioral arena, and to being tethered, and one sham intra-NAc infusion of saline. On the day of the experiment, the infusion cannula was attached via tubing to a 5 uL syringe (Hamilton) mounted on a syringe pump (Harvard apparatus) and threaded through the guide cannula to reach the target structure in each mouse. Mice then received 2.5 ug of bupropion or escitalopram dissolved in 500 nL of saline infused at a rate of 150 nL/min. Mice were allowed to recover for 5-10 min in the home cage after the infusion completed before being placed into the open field arena for 90 min and then were immediately perfused. The brains were processed for cfos immunohistochemistry as described above and imaged as follows. For each of 6-8 sections per mouse, we acquired one image of the NAc^pmSh^ (AP +0.98-1.34 mm from bregma) on the implanted hemisphere (∼300 um x ∼300 um, 40x 1.3 NA objective, Nikon A1 confocal microscope running NIS Elements AR 5.02.00 software). Cfos+ cells and their colocalization with TdTomato and/or eGFP markers were then manually counted.

### Slice electrophysiology

Coronal striatal slices were prepared as previously described^53^. *Drd1a-tdTomato* or *Drd2-eGFP* mice (P60-90) were decapitated following deep isoflurane anesthesia. The brain was quickly removed and placed in ice cold cutting solution (bubbled with 95%O2/5%CO2) consisting (in mM): 228 sucrose, 2.5 KCl, 1.0 NaH2PO4, 8 MgSO4, 26 NaHCO3, 20 glucose and 0.5 CaCl2. The forebrain was isolated, mounted on a cutting platform, and hemisected prior to slicing. Coronal slices (200 µm thick) containing the NAc were cut with a vibratome (Leica VT1200s) and transferred to a holding chamber containing warmed (30 C, bubbled with 95%O2/5%CO2) ASCF consisting (in mM): 119 NaCl. 2.5 KCl, 1 NaH2PO4, 1.3 MgCl2, 26 NaHCO3, 10 glucose and 2.5 CaCl2. Slices were incubated for 45 min and then kept at room temperature for 1-1.5 hrs before being transferred to the fixed recording chamber of an upright microscope (BX50WI, Olympus). Slices were perfused with ACSF (bubbled with 95%O2/5%CO2) and maintained at 30-32 C. D1- and D2- MSNs in the medial shell of posterior NAc slices were identified by epifluorescence. To prevent dialysis of intracellular contents, which can interfere with G-protein signaling cascades of interest in this study, cell-attached voltage-clamp recordings were made using patch electrodes containing (in mM): 146 NaCl, 10 HEPES (pH adjusted to 7.4 with NaOH)^57^. Electrode resistances ranged from 6-8 MΩ. Cell-attached recordings were maintained at a holding potential of −10 mV and resistances (> 1 GΩ) were continually monitored by delivering a 4 mV, 100ms voltage-pulse every 20 s. Cell-attached voltage-clamp recordings were acquired with a MultiClamp 700B (Molecular Devices) and recorded in gap free mode (PClamp 10, Molecular Devices). Signals were filtered at 2 KHz and digitized at 10 KHz (National Instruments BNC-2090 or Digidata 1320A, Axon Instruments). All drugs (Sigma Aldrich) were bath applied and diluted to their final concentrations from concentrated stocks (1000-2000x in dH20). Spontaneous action potentials (APs) were generated by addition of barium chloride (Ba2+, 0.5 mM) and 4-aminopyridine (4-AP, 0.15 mM) to the ACSF. Once steady AP firing had been achieved (∼5 min) slices were exposed to either serotonin (5-HT, 25 µM) or dopamine (DA, 100 µM) for 4 min. Effects on AP frequency were quantitated in a 3 min window before and the last 3 min of drug exposure. APs were analyzed with MiniAnalysis software (Synaptosoft).

### Statistics and reproducibility

Data were analyzed in Graphpad Prism and R. Data that met assumptions of normality and equal variance (assessed by inspection of the fitted-values vs residuals and/or Q-Q plots) were analyzed using parametric tests, with two-way ANOVAs followed by Holm-Sidak or Bonferroni post-hoc tests for two-factor designs and t-tests or one-way ANOVAs followed by Holm-Sidak post-hoc tests for one factor designs. Paired or repeated measures and the Greenhouse-Geisser correction were used where appropriate. Data that did not meet assumptions of normality and equal variance underwent the Box-Cox transformation and parametric tests were then used on the transformed data, or the data were analyzed with nonparametric tests instead (Wilcoxon signed-rank test, Friedman test followed by Dunn’s multiple comparisons tests). All hypothesis tests were two-tailed and data are shown as mean +/− sem.

## ACKNOWLEDGMENTS

We thank members of the Malenka lab, STAAR lab, and Austen B. Casey for discussions and the Stanford Gene Vector and Virus Core for reagents. This work was supported by philanthropic funds donated to the Nancy Pritzker Laboratory at Stanford University, an NSF Graduate Research Fellowship (D.F.C.P.), an HHMI Gilliam Fellowship for Advanced Study (D.F.C.P. and R.C.M.), a Stanford Vice Provost for Undergraduate Education Major Grant (M.Y.G.), a Brain & Behavior Research Foundation Young Investigator Grant (N.E.), a Burroughs Wellcome Fund Career Award for Medical Scientists (N.E.), a Stanford NeuroChoice Initiative Pilot Award (N.E.), a Simons Foundation Bridge to Independence Award (N.E.), a Stanford NeuroChoice Initiative Pilot Award (N.E.), a grant from the Stanford Wu Tsai Neurosciences Institute (R.C.M.), a grant from the UCSF Dolby Family Center for Mood Disorders (R.C.M.), and NIH grants K99DA056573 (M.B.P.), K08MH123791 (N.E.), R01MH138645 (N.E.), and P50DA042012 (R.C.M.).

## AUTHOR CONTRIBUTIONS

D.F.C.P., N.E., and R.C.M. conceived the study. D.F.C.P. designed the experiments with input from all coauthors. M.Y.G collected fluorescence *in situ* hybridization data with assistance from M.B.P. M.Y.G collected cfos data with assistance from M.B.P. W.M. collected electrophysiology data. D.F.C.P. collected presynaptic bouton tracing data. D.F.C.P. analyzed the data with assistance from M.Y.G. and W.M. The manuscript was written by D.F.C.P., N.E., and R.C.M. with input from W.M. and edited by all authors.

## DATA AVAILABILITY

Data in this paper are available from the corresponding author upon request, except for RNAseq data analyzed in Extended Data Fig. 1 which were generated in reference 22 and are available on FigShare at the following link: https://figshare.com/projects/Continuous_and_discrete_neuron_types_of_the_adult_murine_striatum/69080.

## CODE AVAILABILITY

Code used in this study is available from the corresponding author upon request.

## EXTENDED DATA AND FIGURE LEGENDS

**Extended Data Fig. 1:**
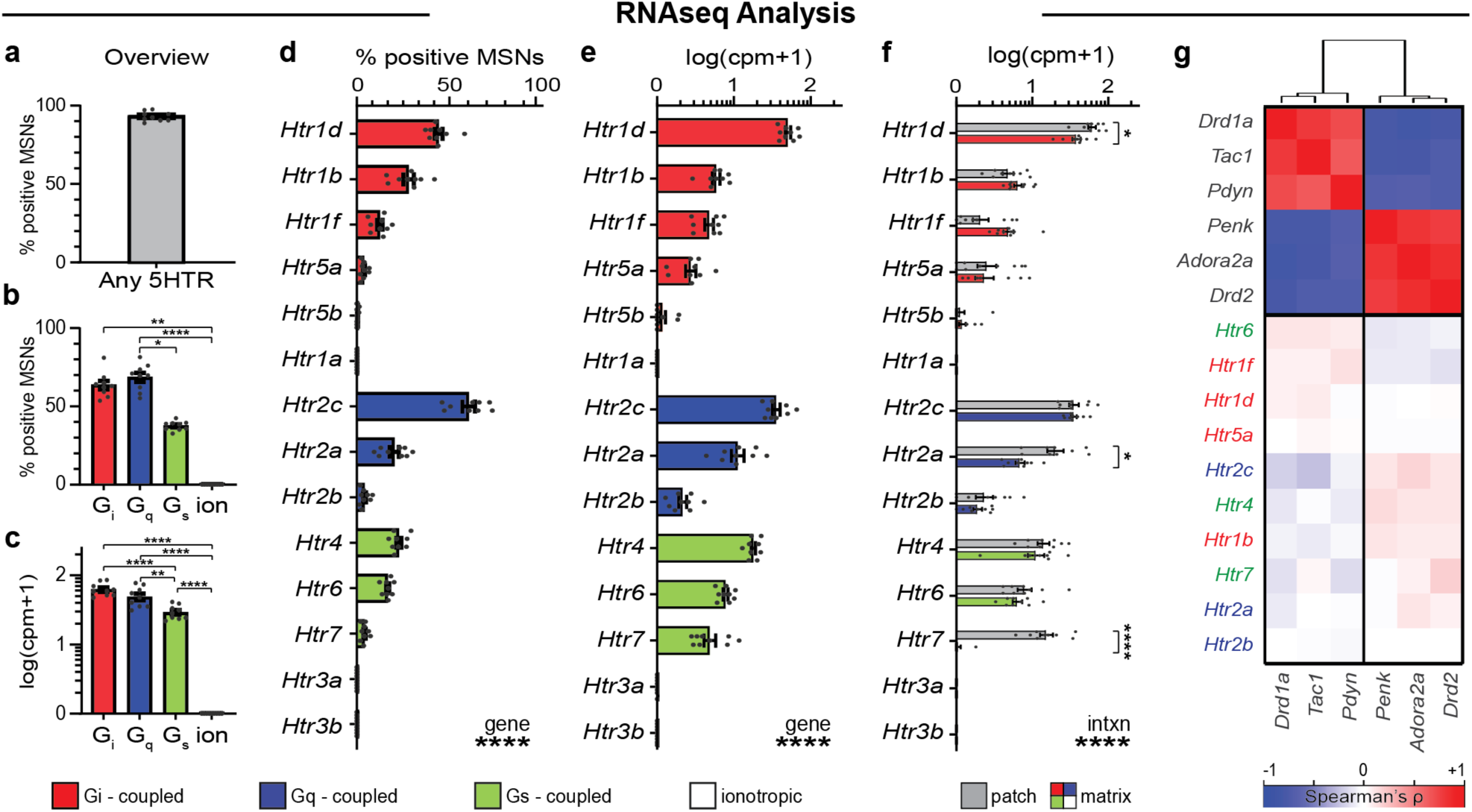
RNAseq analysis of 5HT receptor expression in striatal MSNs. **a,** Percentage of MSNs expressing at least one copy of any 5HT receptor gene. **b,** Same as **a** but broken down by receptor type. **c,** Average expression level of 5HT receptor genes broken down by receptor type. **d,** Percentage of MSNs expressing at least one copy of each 5HT receptor gene. **e,** Average expression level of each 5HT receptor gene. **f,** Same as **e** but broken down by patch and matrix. **g,** Heatmap showing correlation between expression levels of D1 and D2 MSN marker genes and 5HT receptor genes. Data in **a**-**g** are from ref 22. cpm, counts per million reads.

**Extended Data Fig. 2:**
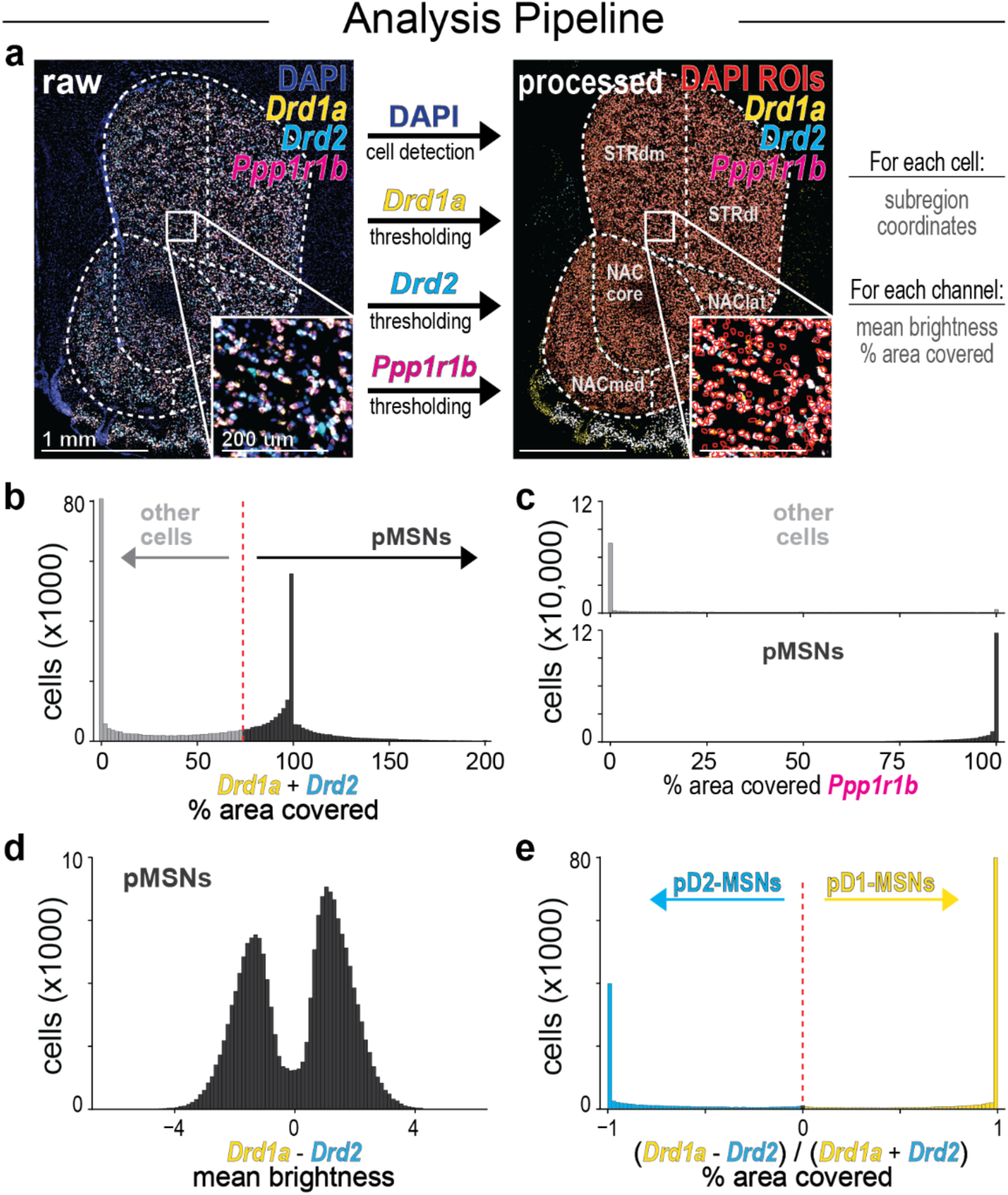
Analysis pipeline for fluorescence *in situ* hybridization experiments. **a,** Depiction of image analysis pipeline for fluorescence *in situ* hybridization experiments. Anatomical regions were manually annotated, cell-detection was done based on the DAPI signal, and for the remaining fluorescence channels the brightness (before thresholding) and fraction of area covered (after thresholding) were measured in each detected cell. **b,** Histogram depicting percentage of area covered by *Drd1a* or *Drd2* signal in each cell. **c,** Histogram depicting percentage of area covered by signal from the MSN marker *Ppp1r1b* in putative non-MSNs (top) and pMSNs (bottom). **d,** Histogram depicting the difference in brightness between the *Drd1a* and *Drd2* signals for all pMSN cells. **e**, Histogram depicting the cell-type index score for all pMSNs, defined on the basis of their *Drd1a* and *Drd2* signals.

**Extended Data Fig. 3:**
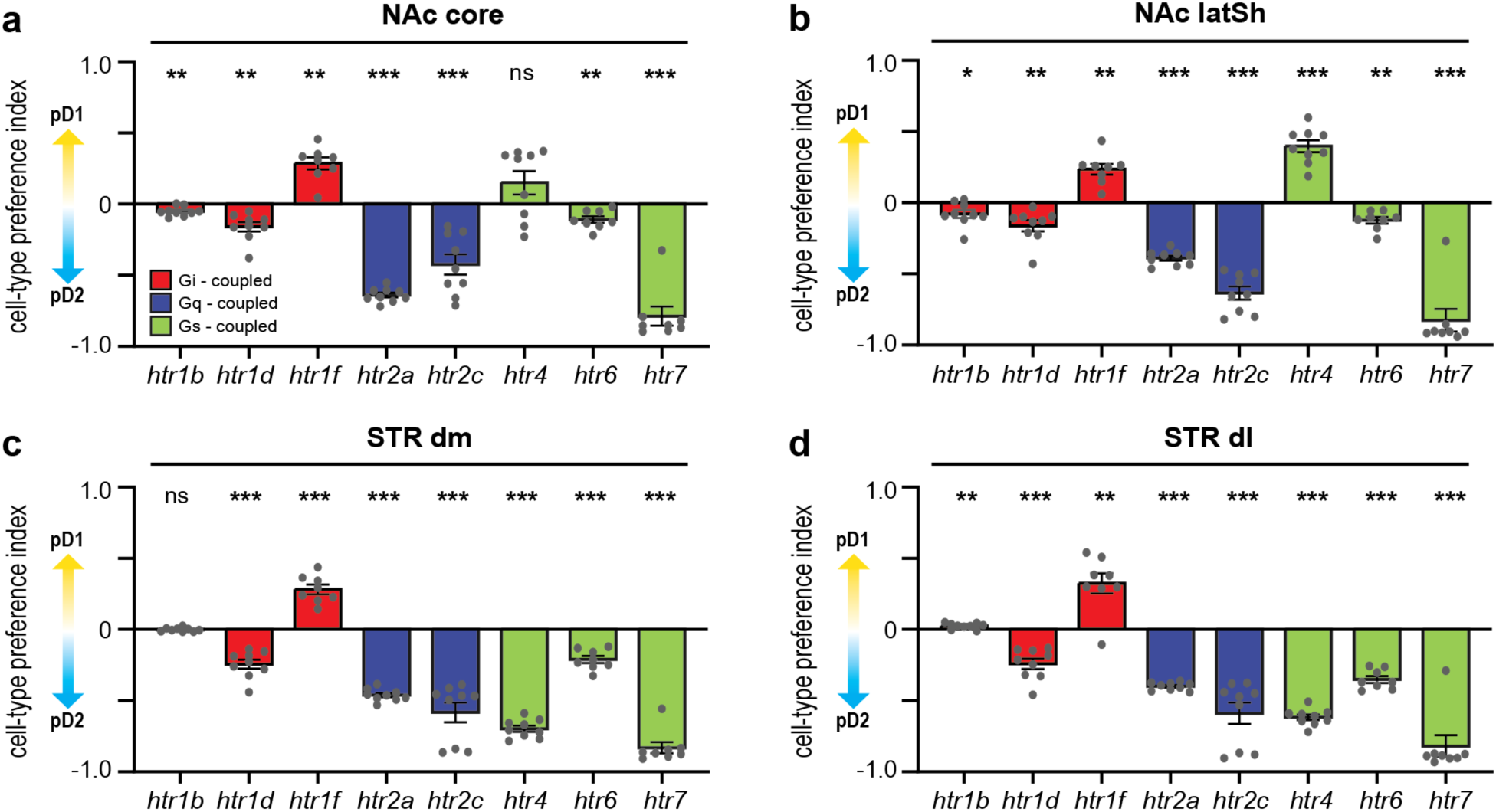
Cell-type preference analysis for 5HT receptor expression across striatal regions. **a,** Cell-type preference analysis for 5HT receptor expression in the NAc^core^. **b,** Same as **a** but for the NAc^latSh^. **c,** Same as **a** but for the STR^dm^. **d,** Same as **a** but for the STR^dl^.

**Extended Data Fig. 4:**
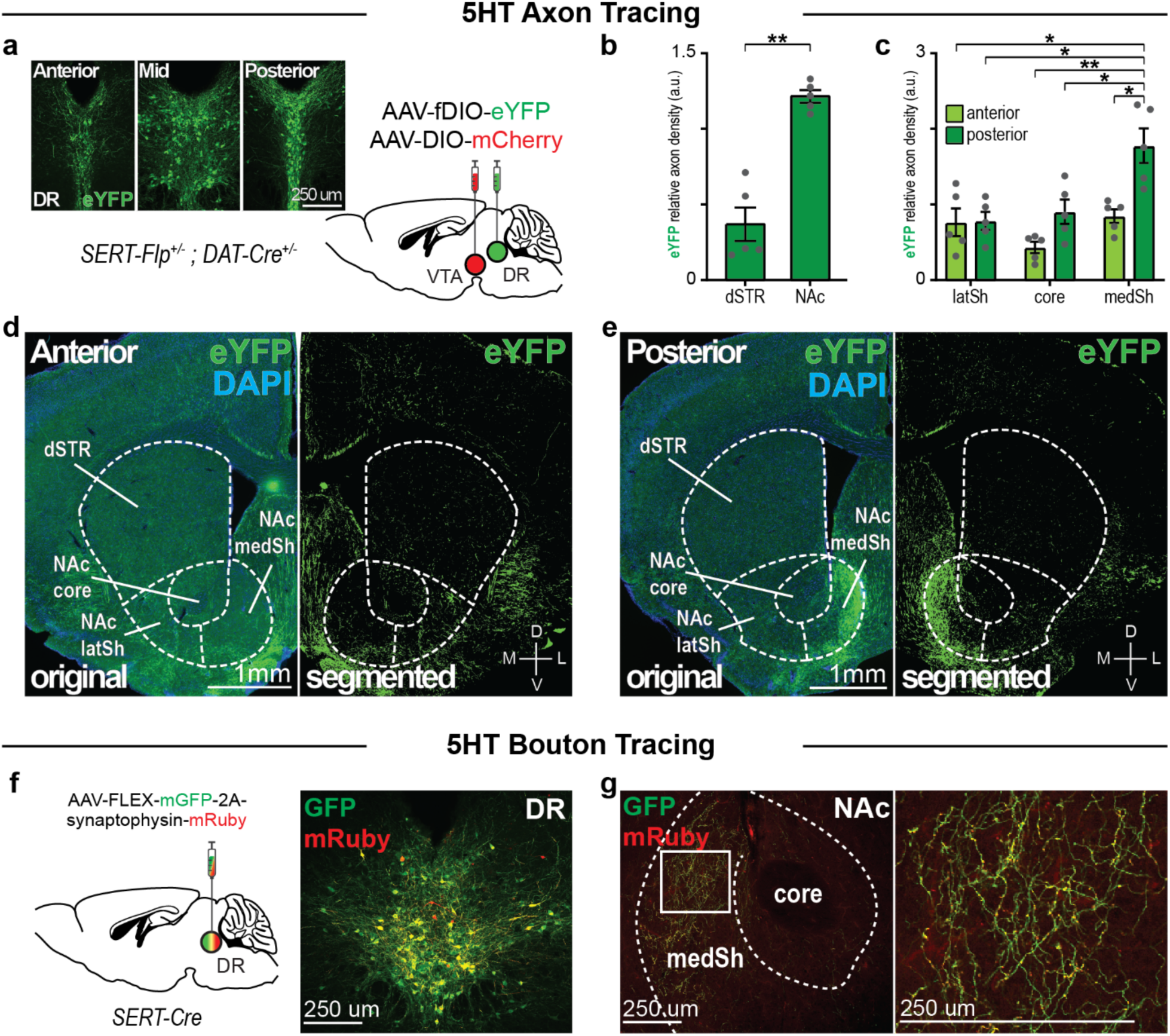
5HT inputs to striatum are topographically organized. **a,** Surgical strategy (right) for labeling DR^5HT^ neurons and example images of the injection site (left). **c,** Quantification of axon density in the dSTR and NAc (left). **c,** Quantification of axon density across subregions of the NAc. **d-e,** Example images of labeled DR^5HT^ in the anterior (**d**) and posterior (**e**) striatum. **f,** Left, surgical strategy for labeling axons and presynaptic boutons in DR^5HT^ neurons. Right, example image of the injection site. **g,** Left, example images of labeled DR^5HT^ axons and presynaptic boutons in the NAc. Right, inset of boxed region on the left.

**Extended Data Fig. 5:**
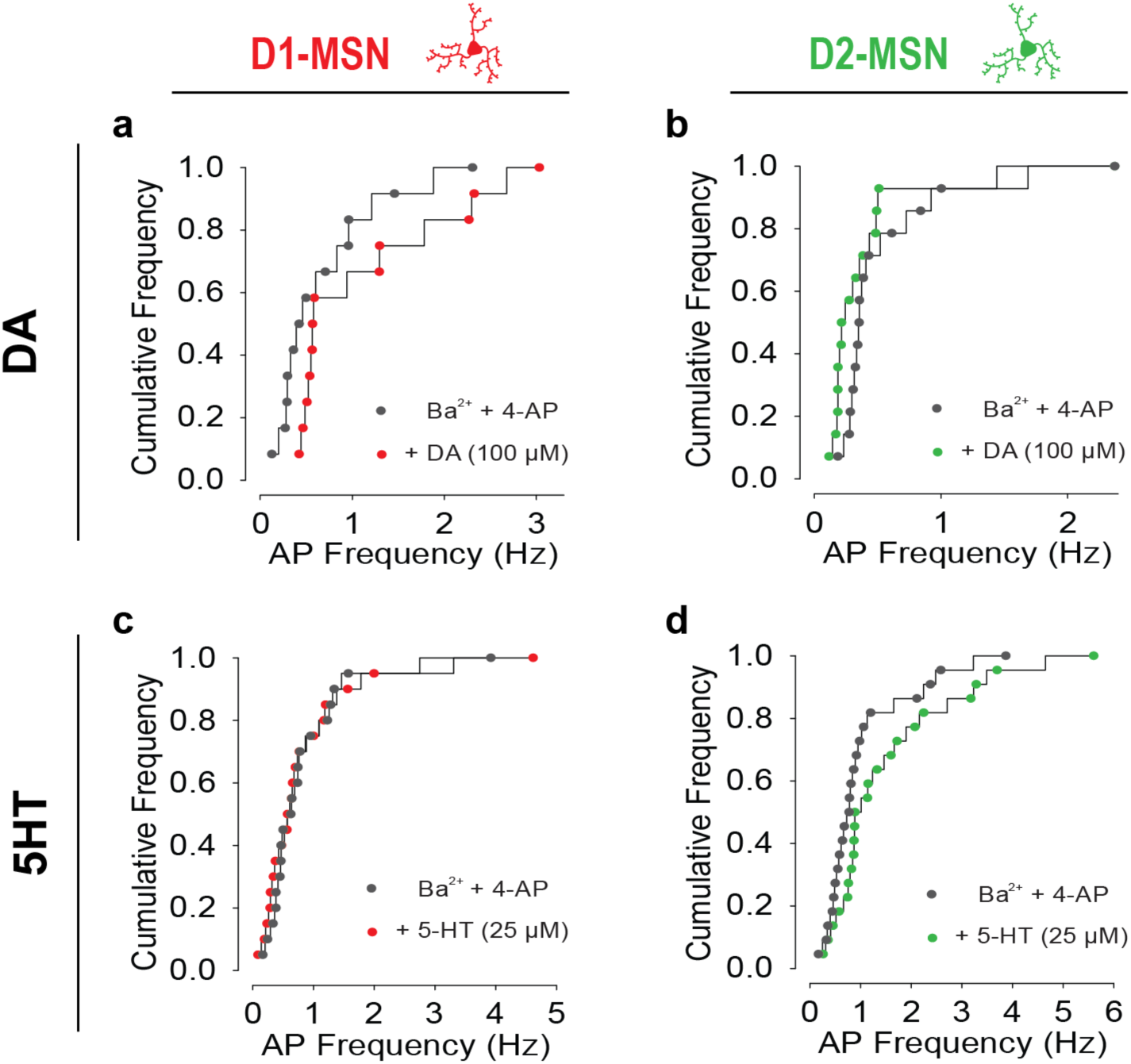
Cumulative frequency plots for cell-attached recording experiments. **a-b,** Cumulative frequency plots for cell-attached recordings from D1- (**a**) and D2- (**b**) MSNs before and after bath application of DA. **c-d,** Same as **a-b** but for bath application of 5HT.

