## Supplemental Table 1 - statistics for "Opponent regulation of striatal output pathways by dopamine and serotonin"

| Figure | Group | n | units | Statistical Tests (all two-tailed) | Test Statistics significant results in red | P vales | Post-hoc Tests (all two-tailed) | | P values significant results in red |
|---|---|---|---|---|---|---|---|---|---|
| 1a | STRdl | 9 | mice | Two-Way RM ANOVA with Holm-Sidak post-hoc tests | | | STRdl | pD1 vs pD2 | 0.0225 |
| | STRdm | 9 | mice | subregion | F (2.017, 16.13) = 114.9 | P<0.0001 | STRdm | pD1 vs pD2 | 0.9372 |
| | NACmed | 9 | mice | cell-type | F (1.000, 8.000) = 1.414 | P=0.2685 | NACmed | pD1 vs pD2 | 0.0018 |
| | NACcor | 9 | mice | subregion x cell-type | F (1.266, 10.13) = 19.71 | P=0.0008 | NACcor | pD1 vs pD2 | 0.0061 |
| | NAClat | 9 | mice | | | | NAClat | pD1 vs pD2 | 0.0395 |
| 1b | STRdl | 9 | mice | Two-Way RM ANOVA with Holm-Sidak post-hoc tests | | | STRdl | pD1 vs pD2 | 0.0004 |
| | STRdm | 9 | mice | subregion | F (2.697, 21.57) = 24.41 | P<0.0001 | STRdm | pD1 vs pD2 | 0.0003 |
| | NACmed | 9 | mice | cell-type | F (1.000, 8.000) = 39.54 | P=0.0002 | NACmed | pD1 vs pD2 | 0.0727 |
| | NACcor | 9 | mice | subregion x cell-type | F (3.165, 25.32) = 27.00 | P<0.0001 | NACcor | pD1 vs pD2 | 0.0008 |
| | NAClat | 9 | mice | | | | NAClat | pD1 vs pD2 | 0.0008 |
| 1c | STRdl | 8 | mice | Two-Way RM ANOVA | | | | pD1 vs pD2 | |
| | STRdm | 8 | mice | subregion | F (1.781, 12.47) = 26.82 | P<0.0001 | | pD1 vs pD2 | |
| | NACmed | 8 | mice | cell-type | F (1.000, 7.000) = 32.87 | P=0.0007 | | pD1 vs pD2 | |
| | NACcor | 8 | mice | subregion x cell-type | F (1.618, 11.33) = 3.708 | P=0.0648 | | pD1 vs pD2 | |
| | NAClat | 8 | mice | | | | | pD1 vs pD2 | |
| 1d | STRdl | 9 | mice | Two-Way RM ANOVA with Holm-Sidak post-hoc tests on Box-Cox transformed data (lambda=0.5) | | | STRdl | pD1 vs pD2 | <0.0001 |
| | STRdm | 9 | mice | subregion | F (2.023, 16.19) = 93.05 | P<0.0001 | STRdm | pD1 vs pD2 | <0.0001 |
| | NACmed | 9 | mice | cell-type | F (1.000, 8.000) = 449.9 | P<0.0001 | NACmed | pD1 vs pD2 | 0.0003 |
| | NACcor | 9 | mice | subregion x cell-type | F (1.820, 14.56) = 17.56 | P=0.0002 | NACcor | pD1 vs pD2 | <0.0001 |
| | NAClat | 9 | mice | | | | NAClat | pD1 vs pD2 | <0.0001 |
| 1e | STRdl | 9 | mice | Two-Way RM ANOVA with Holm-Sidak post-hoc tests on Box-Cox transformed data (lambda=0) | | | STRdl | pD1 vs pD2 | 0.0015 |
| | STRdm | 9 | mice | subregion | F (1.416, 11.33) = 51.94 | P<0.0001 | STRdm | pD1 vs pD2 | 0.0015 |
| | NACmed | 9 | mice | cell-type | F (1.000, 8.000) = 49.89 | P=0.0001 | NACmed | pD1 vs pD2 | <0.0001 |
| | NACcor | 9 | mice | subregion x cell-type | F (1.950, 15.60) = 5.228 | P=0.0189 | NACcor | pD1 vs pD2 | 0.0015 |
| | NAClat | 9 | mice | | | | NAClat | pD1 vs pD2 | <0.0001 |
| 1f | STRdl | 9 | mice | Two-Way RM ANOVA with Holm-Sidak post-hoc tests on Box-Cox transformed data (lambda=0) | | | STRdl | pD1 vs pD2 | <0.0001 |
| | STRdm | 9 | mice | subregion | F (2.396, 19.16) = 259.4 | P<0.0001 | STRdm | pD1 vs pD2 | <0.0001 |
| | NACmed | 9 | mice | cell-type | F (1.000, 8.000) = 21.55 | P=0.0017 | NACmed | pD1 vs pD2 | <0.0001 |
| | NACcor | 9 | mice | subregion x cell-type | F (2.109, 16.87) = 163.2 | P<0.0001 | NACcor | pD1 vs pD2 | 0.4148 |
| | NAClat | 9 | mice | | | | NAClat | pD1 vs pD2 | 0.0001 |
| 1g | STRdl | 8 | mice | Two-Way RM ANOVA | | | STRdl | pD1 vs pD2 | 0.0003 |
| | STRdm | 8 | mice | subregion | F (2.747, 19.23) = 25.91 | P<0.0001 | STRdm | pD1 vs pD2 | 0.0009 |
| | NACmed | 8 | mice | cell-type | F (1.000, 7.000) = 41.96 | P=0.0003 | NACmed | pD1 vs pD2 | 0.1456 |
| | NACcor | 8 | mice | subregion x cell-type | F (2.180, 15.26) = 72.43 | P<0.0001 | NACcor | pD1 vs pD2 | 0.0047 |
| | NAClat | 8 | mice | | | | NAClat | pD1 vs pD2 | 0.0047 |
| 1h | STRdl | 8 | mice | Two-Way RM ANOVA with Holm-Sidak post-hoc tests on Box-Cox transformed data (lambda=0) | | | STRdl | pD1 vs pD2 | 0.0002 |
| | STRdm | 8 | mice | subregion | F (2.339, 16.37) = 23.56 | P<0.0001 | STRdm | pD1 vs pD2 | <0.0001 |
| | NACmed | 8 | mice | cell-type | F (1.000, 7.000) = 92.50 | P<0.0001 | NACmed | pD1 vs pD2 | 0.0002 |
| | NACcor | 8 | mice | subregion x cell-type | F (2.260, 15.82) = 11.53 | P=0.0006 | NACcor | pD1 vs pD2 | 0.0001 |
| | NAClat | 8 | mice | | | | NAClat | pD1 vs pD2 | 0.0002 |
| 1k | *htr1b* | 9 | mice | One-Sample T-tests (hypothetical mean = 0) with Holm's multiplicity adjustment | t=5.072, df=8 | 0.004 | | | |
| | *htr1d* | 9 | mice | | t=1.731, df=8 | 0.2436 | | | |
| | *htr1f* | 8 | mice | | t=3.583, df=7 | 0.0267 | | | |
| | *htr2a* | 9 | mice | | t=5.577, df=8 | 0.0025 | | | |
| | *htr2c* | 9 | mice | | t=14.99, df=8 | 0.0008 | | | |
| | *htr4* | 9 | mice | | t=26.56, df=8 | 0.0008 | | | |
| | *htr6* | 8 | mice | | t=1.372, df=7 | 0.2436 | | | |
| | *htr7* | 8 | mice | | t=8.974, df=7 | 0.0008 | | | |
| 2c | pD1-MSNs | 32 cells / 10 mice | | Two-Way RM ANOVA with Holm-Sidak post-hoc tests | | | pD1-MSNs | ACSF vs BA/4AP | <0.0001 |
| | pD2-MSNs | 36 cells / 13 mice | | cell-type | F (1, 66) = 0.08219 | P=0.7753 | pD2-MSNs | ACSF vs BA/4AP | <0.0001 |
| | | | | bath solution | F (1, 66) = 0.08219 | P=0.7753 | | | |
| | | | | cell-type x bath solution | F (1, 66) = 77.52 | P<0.0001 | | | |
| 2e | pD1-MSNS | 12 cells / 4 mice | | Wilcoxon matched-pairs signed rank test | | 0.0015 | | | |
| 2g | pD1-MSNS | 14 cells / 4 mice | | Wilcoxon matched-pairs signed rank test | | 0.0256 | | | |
| 2i | pD2-MSNS | 20 cells / 6 mice | | Wilcoxon matched-pairs signed rank test | | 0.6477 | | | |
| 2k | pD2-MSNS | 22 cells / 9 mice | | Wilcoxon matched-pairs signed rank test | | 0.0006 | | | |
| 2l | pD1-MSN DA | 12 cells / 4 mice | | Two-Way ANOVA with Holm-Sidak post-hoc tests | | | DA | pD1 vs pD2 | 0.0056 |
| | pD2-MSN DA | 14 cells / 4 mice | | cell-type x bath solution | F (1, 64) = 17.58 | P<0.0001 | 5HT | pD1 vs pD2 | 0.0051 |
| | pD1-MSN 5HT | 20 cells / 6 mice | | bath solution | F (1, 64) = 0.9719 | P=0.3279 | | | |
| | pD2-MSN 5HT | 22 cells / 9 mice | | cell-type | F (1, 64) = 0.09852 | P=0.7546 | | | |
| 2m | pD1-MSN DA | 12 cells / 4 mice | | One-Sample T-tests (hypothetical mean = 1) with Holm's multiplicity adjustment | t=3.968, df=11 | 0.0066 | | | |
| | pD2-MSN DA | 14 cells / 4 mice | | | t=2.712, df=13 | 0.0356 | | | |
| | pD1-MSN 5HT | 20 cells / 6 mice | | | t=0.7315, df=19 | 0.4734 | | | |
| | pD2-MSN 5HT | 22 cells / 9 mice | | | t=3.670, df=21 | 0.0056 | | | |
| 2q (left) | BUP | 5 | mice | Unpaired T-test | t(8)=2.440 | 0.0406 | | | |
| | ESC | 5 | mice | | | | | | |
| 2q (right) | pD1-MSN BUP | 5 | mice | Two-Way RM ANOVA with Holm-Sidak post-hoc tests | | | BUP | pD1 vs pD2 | 0.0026 |
| | pD2-MSN BUP | 5 | mice | drug x cell-type | F (1, 8) = 45.10 | P=0.0002 | ESC | pD1 vs pD2 | 0.0017 |
| | pD1-MSN ESC | 5 | mice | drug | F (1, 8) = 1.000 | P=0.3466 | | | |
| | pD2-MSN ESC | 5 | mice | cell-type | F (1, 8) = 0.3866 | P=0.5514 | | | |
| ED1b | Gi | 9 | mice | Friedman Test followed by Dunn's post-hoc test | 25.11 | <0.0001 | | Gi vs. Gq | >0.9999 |
| | Gq | 9 | mice | | | | | Gi vs. Gs | 0.2146 |
| | Gs | 9 | mice | | | | | Gi vs. Ion | 0.0011 |
| | ion | 9 | mice | | | | | Gq vs. Gs | 0.0279 |
| | | | | | | | | Gq vs. Ion | <0.0001 |
| | | | | | | | | Gs vs. Ion | 0.6021 |
| ED1c | Gi | 9 | mice | One-Way RM ANOVA with Holm-Sidak's post-hoc tests | F (1.890, 15.12) = 714.6 | P<0.0001 | | Gi vs. Gq | 0.113 |
| | Gq | 9 | mice | | | | | Gi vs. Gs | <0.0001 |
| | Gs | 9 | mice | | | | | Gi vs. Ion | <0.0001 |
| | ion | 9 | mice | | | | | Gq vs. Gs | 0.0051 |
| | | | | | | | | Gq vs. Ion | <0.0001 |
| | | | | | | | | Gs vs. Ion | <0.0001 |
| ED1d | Htr1d | 9 | mice | Friedman Test | 112.6 | <0.0001 | | | |
| | Htr1b | 9 | mice | | | | | | |
| | Htr1f | 9 | mice | | | | | | |
| | Htr5a | 9 | mice | | | | | | |
| | Htr5b | 9 | mice | | | | | | |
| | Htr1a | 9 | mice | | | | | | |
| | Htr2c | 9 | mice | | | | | | |
| | Htr2a | 9 | mice | | | | | | |
| | Htr2b | 9 | mice | | | | | | |
| | Htr4 | 9 | mice | | | | | | |
| | Htr6 | 9 | mice | | | | | | |
| | Htr7 | 9 | mice | | | | | | |
| | Htr3a | 9 | mice | | | | | | |
| | Htr3b | 9 | mice | | | | | | |
| ED1e | Htr1d | 9 | mice | Friedman Test | 111.8 | <0.0001 | | | |
| | Htr1b | 9 | mice | | | | | | |
| | Htr1f | 9 | mice | | | | | | |
| | Htr5a | 9 | mice | | | | | | |
| | Htr5b | 9 | mice | | | | | | |
| | Htr1a | 9 | mice | | | | | | |
| | Htr2c | 9 | mice | | | | | | |
| | Htr2a | 9 | mice | | | | | | |
| | Htr2b | 9 | mice | | | | | | |
| | Htr4 | 9 | mice | | | | | | |
| | Htr6 | 9 | mice | | | | | | |
| | Htr7 | 9 | mice | | | | | | |
| | Htr3a | 9 | mice | | | | | | |
| | Htr3b | 9 | mice | | | | | | |
| ED1f | Htr1d | 9 | mice | Two-Way RM ANOVA with Bonferroni's post-hoc tests | | | Htr1d | D1 vs D2 | 0.0451 |
| | Htr1b | 9 | mice | gene | F (4.577, 36.62) = 136.5 | P<0.0001 | Htr1b | D1 vs D2 | >0.9999 |
| | Htr1f | 9 | mice | striatal compartment | F (1.000, 8.000) = 31.14 | P=0.0005 | Htr1f | D1 vs D2 | 0.371 |
| | Htr5a | 9 | mice | gene x striatal compartment | F (3.480, 27.84) = 11.97 | P<0.0001 | Htr5a | D1 vs D2 | >0.9999 |
| | Htr5b | 9 | mice | | | | Htr5b | D1 vs D2 | >0.9999 |
| | Htr1a | 9 | mice | | | | Htr1a | D1 vs D2 | n/a, all samples are 0 |
| | Htr2c | 9 | mice | | | | Htr2c | D1 vs D2 | >0.9999 |
| | Htr2a | 9 | mice | | | | Htr2a | D1 vs D2 | 0.0203 |
| | Htr2b | 9 | mice | | | | Htr2b | D1 vs D2 | >0.9999 |
| | Htr4 | 9 | mice | | | | Htr4 | D1 vs D2 | >0.9999 |

| | | | | | | | | | |
|---|---|---|---|---|---|---|---|---|---|
| | Htr6 | 9 | mice | | | | Htr6 | D1 vs D2 | >0.9999 |
| | Htr7 | 9 | mice | | | | Htr7 | D1 vs D2 | <0.0001 |
| | Htr3a | 9 | mice | | | | Htr3a | D1 vs D2 | n/a, all samples are 0 |
| | Htr3b | 9 | mice | | | | Htr3b | D1 vs D2 | n/a, all samples are 0 |
| ED3a | *htr1b* | 9 | mice | One-Sample T-tests (hypothetical mean = 0) with Holm's multiplicity adjustment | t=4.639, df=8 | 0.0051 | | | |
| | *htr1d* | 9 | mice | | t=4.915, df=8 | 0.0048 | | | |
| | *htr1f* | 8 | mice | | t=6.733, df=7 | 0.0018 | | | |
| | *htr2a* | 9 | mice | | t=38.54, df=8 | 0.0008 | | | |
| | *htr2c* | 9 | mice | | t=6.015, df=8 | 0.0018 | | | |
| | *htr4* | 9 | mice | | t=1.819, df=8 | 0.1064 | | | |
| | *htr6* | 8 | mice | | t=4.651, df=7 | 0.0051 | | | |
| | *htr7* | 8 | mice | | t=11.77, df=7 | 0.0008 | | | |
| ED3b | *htr1b* | 9 | mice | One-Sample T-tests (hypothetical mean = 0) with Holm's multiplicity adjustment | t=2.803, df=8 | 0.0231 | | | |
| | *htr1d* | 9 | mice | | t=4.160, df=8 | 0.0064 | | | |
| | *htr1f* | 8 | mice | | t=6.165, df=7 | 0.002 | | | |
| | *htr2a* | 9 | mice | | t=22.19, df=8 | 0.0008 | | | |
| | *htr2c* | 9 | mice | | t=14.02, df=8 | 0.0008 | | | |
| | *htr4* | 9 | mice | | t=9.544, df=8 | 0.0008 | | | |
| | *htr6* | 8 | mice | | t=5.293, df=7 | 0.0033 | | | |
| | *htr7* | 8 | mice | | t=10.33, df=7 | 0.0008 | | | |
| ED3c | *htr1b* | 9 | mice | One-Sample T-tests (hypothetical mean = 0) with Holm's multiplicity adjustment | t=0.2304, df=8 | 0.8236 | | | |
| | *htr1d* | 9 | mice | | t=7.646, df=8 | 0.0008 | | | |
| | *htr1f* | 8 | mice | | t=8.281, df=7 | 0.0008 | | | |
| | *htr2a* | 9 | mice | | t=32.16, df=8 | 0.0008 | | | |
| | *htr2c* | 9 | mice | | t=8.478, df=8 | 0.0008 | | | |
| | *htr4* | 9 | mice | | t=32.38, df=8 | 0.0008 | | | |
| | *htr6* | 8 | mice | | t=8.519, df=7 | 0.0008 | | | |
| | *htr7* | 8 | mice | | t=20.73, df=7 | 0.0008 | | | |
| ED3d | *htr1b* | 9 | mice | One-Sample T-tests (hypothetical mean = 0) with Holm's multiplicity adjustment | t=3.602, df=8 | 0.007 | | | |
| | *htr1d* | 9 | mice | | t=6.573, df=8 | 0.0008 | | | |
| | *htr1f* | 8 | mice | | t=4.626, df=7 | 0.0048 | | | |
| | *htr2a* | 9 | mice | | t=42.66, df=8 | 0.0008 | | | |
| | *htr2c* | 9 | mice | | t=7.830, df=8 | 0.0008 | | | |
| | *htr4* | 9 | mice | | t=31.31, df=8 | 0.0008 | | | |
| | *htr6* | 8 | mice | | t=14.28, df=7 | 0.0008 | | | |
| | *htr7* | 8 | mice | | t=10.79, df=7 | 0.0008 | | | |
| ED4b | dSTR | 5 | mice | Paired T-test | t(4)=5.668 | 0.0048 | | | |
| | Nac | 5 | mice | | | | | | |
| ED4c | latSh | 5 | mice | Two-Way RM ANOVA with Holm-Sidak post-hoc tests | | | medSh:Posterior | latSh:Anterior | 0.0151 |
| | core | 5 | mice | subregion | F (2, 8) = 9.525 | P=0.0077 | | latSh:Posterior | 0.0156 |
| | medSh | 5 | mice | AP | F (1, 4) = 9.246 | P=0.0384 | | core:Anterior | 0.0025 |
| | | | | subregion x AP | F (2, 8) = 5.026 | P=0.0386 | | core:Posterior | 0.0288 |
| | | | | | | | | medSh:Anterior | 0.0217 |
| | | | | | | | all other comparisons | | > 0.05 |
